# Redesigning the Eterna100 RNA design benchmark for the Vienna 2 folding engine

**DOI:** 10.1101/2021.08.26.457839

**Authors:** Rohan V. Koodli, Boris Rudolfs, Jonathan Romano, Hannah K. Wayment-Steele, William A. Dunlap, Eterna Participants, Rhiju Das

## Abstract

The rational design of RNA is becoming important for rapidly developing technologies in medicine and biochemistry, spurring development of numerous RNA secondary structure design algorithms and benchmarks to evaluate their performance. However, the problem of RNA design is dependent upon the inverse problem of RNA structure prediction through “folding engines” that predict structure from sequence. We hypothesized that differences in RNA folding engines could impact design algorithms, and recruited an online community of RNA design experts to modify the widely used RNA secondary structure design benchmark Eterna100 to address target unsolvability when changing the folding engine used (ViennaRNA 1 updated to 2). We tested this new Eterna100-V2 benchmark with six RNA design algorithms, and found that the performance of RNA design algorithms changed depending on which folding engine was used in training or evaluation. We further explored algorithms reported to have solved the original Eterna100 and found that incorrect folding engine settings were used in their prior evaluation. As a result, currently, both the original Eterna100-V1 and the modernized Eterna100-V2 benchmark remain unsolved with their intended folding engine settings. This work demonstrates that the design difficulty of RNA structures is intrinsically linked to thermodynamic models and suggests that future RNA design algorithms that can adapt to updated thermodynamic models may result in improved performance. Eterna100-V1 and Eterna100-V2 benchmarks and example solutions are freely available at https://github.com/eternagame/eterna100-benchmarking.

**Author Summary:** Designing RNA molecules that fold to a desired target structure is an algorithmic problem gathering increasing attention due to the emergence of RNA-based therapies and the need for rational design of RNAs. The Eterna100 dataset, a collection of target structures with increasing design difficulty, was designed and selected by players of the crowdsourced RNA game Eterna and has been widely used to benchmark RNA design algorithms. However, these puzzles were originally developed using the now deprecated version 1 of the ViennaRNA folding engine. In this manuscript, we introduce an updated benchmark, called the *Eterna100-V2*. We found that 19 puzzles using Vienna 1 could not be solved in Vienna 2 but that Eterna participants were able to redesign the puzzles with minimal modifications to make them solvable in Vienna 2. We confirmed that the rankings of six RNA design algorithms remained largely consistent between Eterna100-V1 and -V2. No algorithm has solved all the puzzles in either benchmark; several published algorithmic solutions were generated with folding engines run with settings not intended for the Eterna100. Discrepancies in performance from algorithms that rely on folding-engine-specific heuristics suggest that algorithms will benefit from being agnostic to thermodynamic models as these models continue to improve.

## Introduction

RNA has significantly expanded past its original proposed role as an intermediate in the genetic code and as a catalytic scaffold for protein synthesis. RNA has been observed to act as a genetic expression regulator [1], perform catalysis [2], be a scaffold for complex formation [3,4], and be used as a guide by several ribonucleoprotein complexes [5–7]. This increased appreciation for the versatile activity of RNA has led to the recent development of several RNA therapies that include the control of pre-mRNA splicing [8], gene editing and expression [6], and aptamers for binding and sequestering target molecules [9]. It is increasingly understood that RNAs obtain some of their function via secondary structures determined by Watson-Crick-Franklin base-pairing. Structured synthetic RNAs underlie methods to regulate gene expression in bacteria [10] and in eukaryotes [11], sense analytes [12], enhance gene editing systems via synthetic CRISPR guide RNAs [13], and design RNA therapeutics with increased stability [14].

The RNA secondary structure design problem, also known as the inverse folding problem, involves designing an RNA sequence that folds into a target secondary structure as its minimum free energy (MFE) structure given an energy function [15]. Classic RNA inverse folding algorithms used cost function minimization through adaptive random walk [16], structure decomposition [17], minimization of the ensemble defect [18], or a genetic algorithm [19]. Performance of these older algorithms was not well characterized, as benchmarking occurred internally and was performed on well-characterized biological RNA or computationally predicted secondary structures from RNA sequences without selection for complexity or difficulty [17,20,21]. Furthermore, as RNA length and complexity increases, the number of asymmetric and symmetric elements increases, thereby increasing the difficulty of sequence design for these molecules [22].

To address the need for a community-wide standard benchmark for RNA design, Anderson-Lee et al. developed a set of 100 secondary structures published on the Eterna website using RNAfold 1.8.5 [16] from the ViennaRNA package (henceforth referred to as “Vienna 1”; see also **Supplemental Data Table S1**). This Eterna100 benchmark (henceforth referred to as Eterna100-V1) was chosen to showcase secondary structure motifs that were identified as being difficult to design, and the best-performing algorithm at the time [19] solved 54/100. Since the benchmark was published, several algorithms have surpassed this mark using convolutional neural networks [23], human design priors [24], reinforcement learning [25], or a nested Monte Carlo search [26]. However, algorithms have been inconsistent in which secondary structure prediction algorithm, or “folding engine”, they use. For example, EternaBrain [23] used Vienna 1 but Meta-LEARNA [25] used updated thermodynamic parameters in a later ViennaRNA package (ViennaRNA RNAfold 2.1.9 or later versions that we have checked to have byte-identical outputs; henceforth referred to as “Vienna 2”). In addition, two studies which claimed to solve all 100 problems [27,28] of the Eterna100 benchmark with Vienna 1 settings did not use the original Vienna 1 package, but instead emulated Vienna1-like thermodynamic parameters with the Vienna 2 package.

Despite potential confounding of which folding engine is used to train vs. evaluate design algorithms, there has been no systematic evaluation of the effect of folding engines in the training and performance of design algorithms. Here, we challenged Eterna participants to determine if all the puzzles in the original Eterna100 could still be solved using the Vienna 2 set of ViennaRNA parameters. Indeed, participants identified 19 of the 100 structures that were unsolvable in Vienna 2. We then challenged the community to adapt these secondary structures to the Vienna 2 parameter set using a minimal number of insertions and deletions, resulting in the Eterna100-V2 benchmark. We discuss key structural motifs that are intractable in one set of thermodynamic parameters but solvable in the other. We evaluated several state-of-the-art inverse folding algorithms and determined that while their relative performance is similar, methods relying on heuristics specific to Vienna 1 – notably the NEMO method – are less performant on the newer Eterna100-V2 with the Vienna 2 folding engine. The folding engine dependence of algorithms also led us to re-investigate reported solutions to the original Eterna100, and we discovered that prior studies have not emulated the intended Vienna 1 settings, leading to overestimates of algorithm performance. These results show that consideration of folding engine is important in development and evaluation of inverse folding algorithms and that both the original and modernized Eterna100 benchmarks remain unsolved.

## Results

### Thermodynamic differences in motifs result in puzzle unsolvability

Studies developing algorithms like the reinforcement learning approach of Eastman et al. [29], LEARNA [25], SamplingDesign [30], SAMFEO [31], m2dRNAs [32], and FastDesign [33] have used the Eterna100 puzzles as a benchmark, but with the modernized Vienna 2 folding engine instead of the originally intended Vienna 1. To understand whether these studies were correctly evaluating their design method’s performance, we crowdsourced the solution of Eterna100 puzzles with Vienna 2, with the hypothesis that many of the puzzles, which targeted idiosyncrasies of Vienna 1, would not be solvable in the newer folding engine. Eterna participants deemed only 81 secondary structures of the original 100 to be ‘solvable’ in Vienna 2 (**Table 1**, bottom row). We verified that the stochastic algorithm NEMO [26], which is currently state-of-the-art in the Eterna100 with the original Vienna 1 folding engine, was also unable to find solutions within 24-hour timeframes for the remaining 19 problems with the Vienna 2 folding engine (**Table 1**).

**Table 1.** Number of solutions of the Eterna100-V1 and Eterna100-V2 benchmarks by six automated algorithms and human players of Eterna, under two folding engines (Vienna 1 and Vienna 2). Highlighted in bold are the two configurations – Eterna100-V1 in Vienna 1, Eterna100-V2 in Vienna 2 – that reflect the original and modernized benchmark, respectively.

|  | **Eterna100-V1** |  | **Eterna100-V2** |  |
|---|---|---|---|---|
|  | **Vienna 1** | **Vienna 2** | **Vienna 1** | **Vienna 2** |
| **RNAinverse** | 18 | 20 | 19 | 20 |
| **EternaBrain** | 60 | 49 | 65 | 49 |
| **LEARNA** | 67 | 66 | 72 | 69 |
| **SentRNA** | 69 | 63 | 72 | 68 |
| **NEMO** | 92 | 77 | 93 | 86 |
| **DesiRNA** | 90 | 78 | 93 | 92 |
| **Any automated method** | 95 | 78 | 96 | 93 |
| **Eterna (humans)** | **100** | 81 | 99 | **100** |

These 19 puzzles varied in both length and relative complexity, but shared several structural features that gave rise to this discrepancy (**Fig. 1a**). The change in the relative free energies of internal and stem loops between Vienna 1 and Vienna 2 led to several structures with isolated base pairs no longer being solvable in the Vienna 2 parameters. Multi-helix junctions were also found to have lower free energies in Vienna 2 than Vienna 1, which significantly altered puzzles that feature these such as “Kyurem 5” and “Multilooping 6” (**Fig. 1b**).

**Fig 1.**
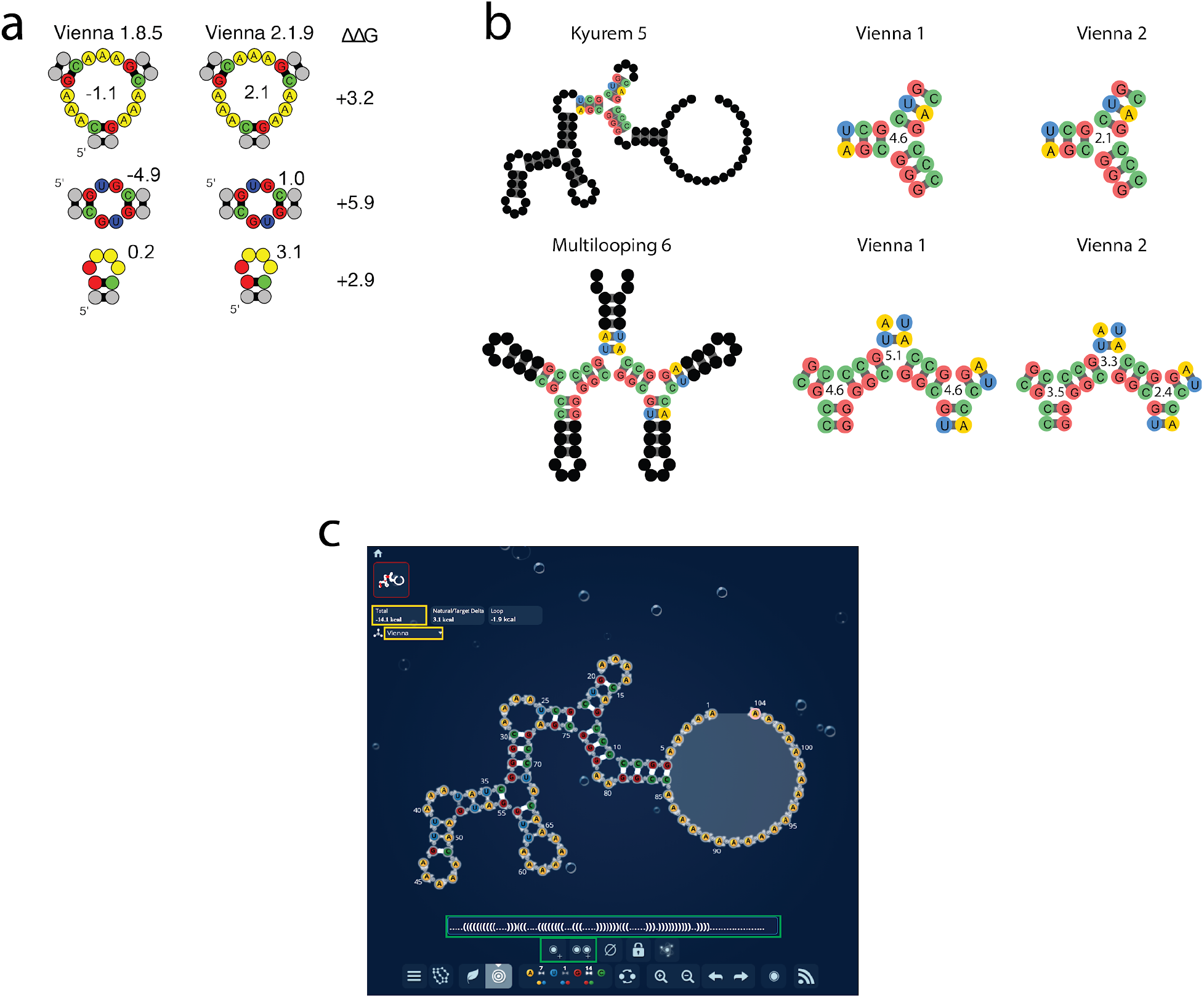
(a)Example free energy differences (ΔΔG, kcal/mol) between identical structures in Vienna 1 and Vienna 2. **(b)** Puzzle-specific differences in free energies of structures. The internal tri-junction in Kyurem 5 has a more stable free energy in Vienna 2. Similar repeated tri-junctions are highlighted in Multilooping 6, with similar changes in free energy from Vienna 1 to 2. **(c)** A screenshot of Eterna’s “Puzzlemaker” interface. Players can see the free energy and folding engine in the top left corner (yellow rectangles). Players can add bases, add base pairs, and interact with the dot-bracket representation of the structure directly (green rectangles).

Large internal loops increased in free energy in Vienna 2 as compared to Vienna 1, which made puzzles containing these, such as “[RNA] Repetitious Sequences 8/10” unsolvable. For example, EternaBrain’s solution to [RNA] Repetitious Sequences 8/10’s Eterna100-V1 structure creates two large internal loops both with free energies of −1.1 kcal/mol. However, if the same structure and solution are used in Vienna 2, the free energies of both internal loops increase to 2.1 kcal/mol (**Fig. 1a, top panels**).

These results motivated us to develop a new set of puzzles for these 19 problems that would comprise a new Eterna100-V2 benchmark to be used with the Vienna 2 folding engine, which has largely displaced the Vienna 1 folding engine for wide use. We solicited these new problems from Eterna participants with a news post (**Supp. Fig. S1**) and provided the Vienna 2 folding engine in the Eterna “puzzlemaker” interface (**Fig. 1c**). These puzzles and example solutions are compiled in **Supplemental Data Table S1** as well as an online repository at https://github.com/eternagame/eterna100-benchmarking.

### Eterna participants redesign target structures with minimal alterations

Given that the original Eterna100-V1 benchmark was designed by players, we hypothesized that players might also be able to design modified structures for these puzzles using a minimum number of structure modifications, while maintaining the constraints of the Eterna software platform. Strategies included adding or removing base pairs to maintain the overall puzzle topology, reducing the size of large loops to mitigate Vienna 2’s increased penalty on large loops. We compared Eterna participant-designed solutions to a “simple” redesign method: taking a set of known player puzzle solutions in Vienna 1 and calculating their minimum free energy structures in Vienna 2. However, these structures differed substantially in shape from the original Eterna100 targets, as quantified by tree edit distance in RNAdistance [16] (**Fig. 2**), and lacked their characteristically difficult features. In all 19 puzzles, this calculated difference compared to the V1 structures was much larger than the difference in the secondary structures that players created in parallel. Some of the most structurally complex puzzles required minimal alterations. “Shooting Star” consists of several multi-helix junctions and long helices that contain 29 isolated base pairs and was made solvable through only 3 base pair additions (**Fig. 3, top**). “Teslagon” consists of a series of loops around a core 7-way junction and was made solvable through the deletion of a single internal loop base (**Fig. 3, middle**). However, some puzzles like “Gladius” required more extensive changes, including breaking structural symmetry, to become solvable in Vienna 2 (**Fig. 3, bottom**).

**Fig 2.**
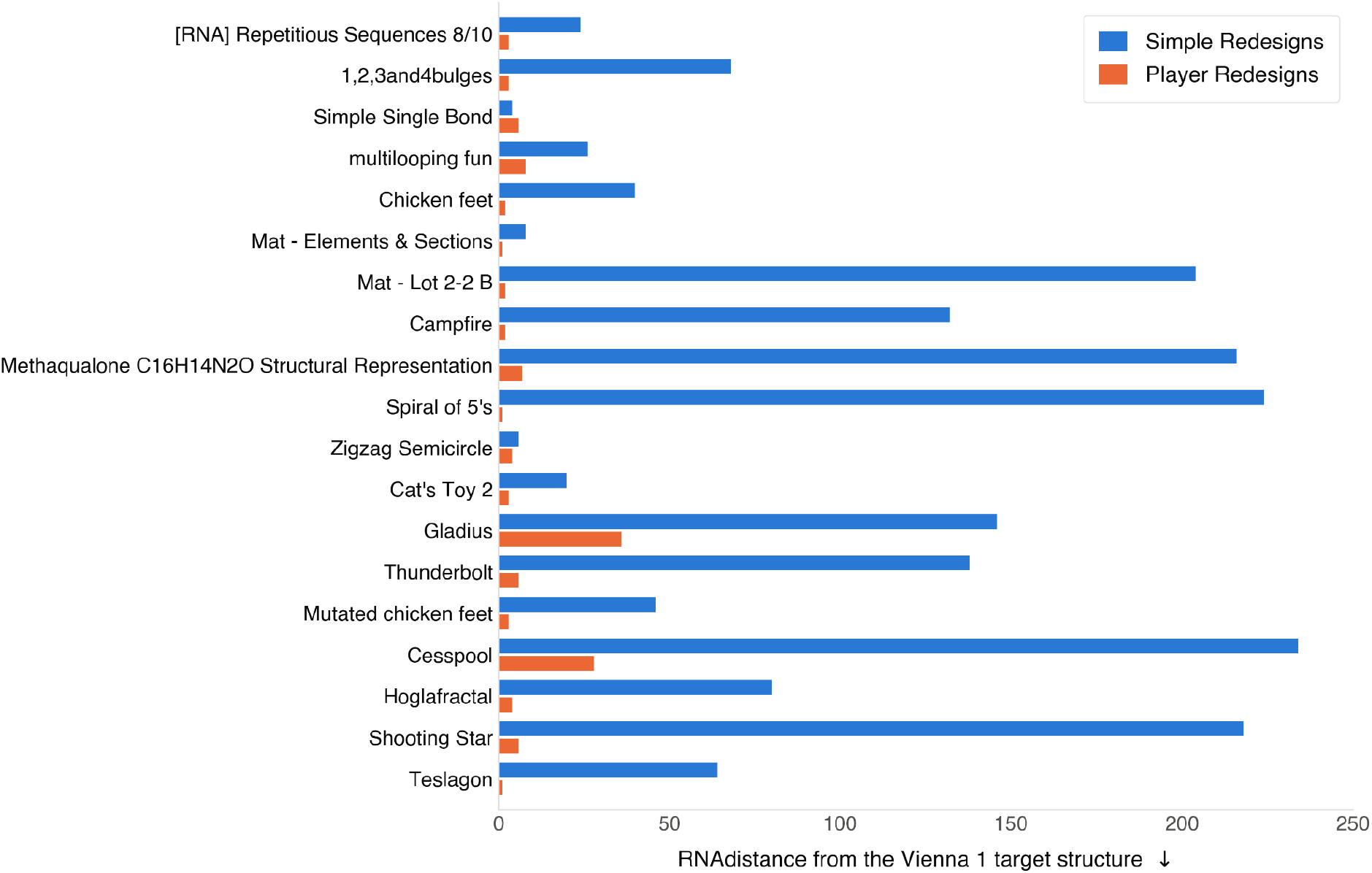
Eterna participant target structure re-designs generally result in fewer structure modifications than a naive re-design approach, quantified by RNAdistance [16].

**Fig 3.**
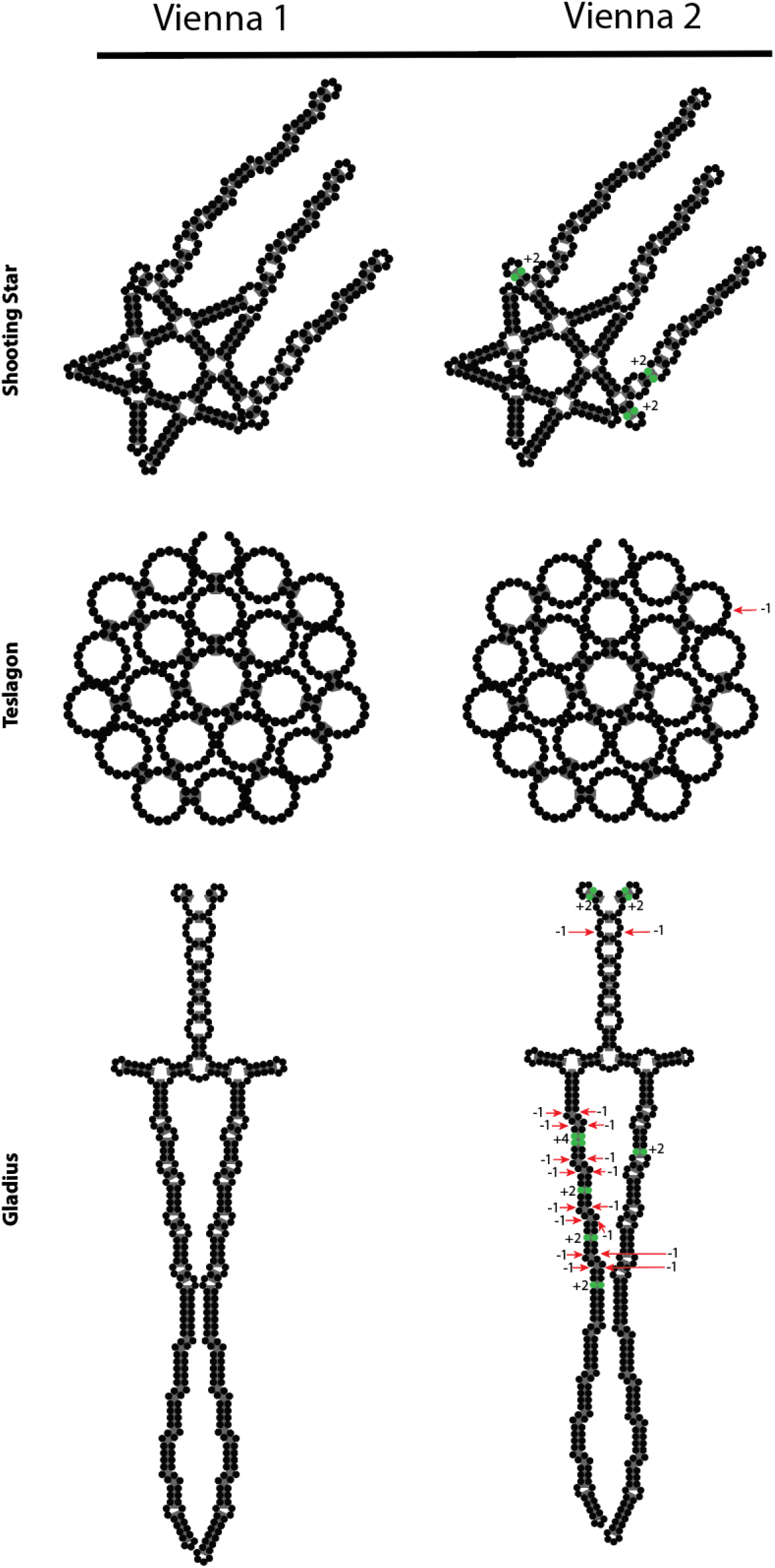
Base pair changes in “Shooting Star” from Vienna 1 to Vienna 2. The green nucleotides indicate the bases that were added to make the structure stable in Vienna 2, and the red indicate deletions.

### Redesigned target structures are largely still solvable using Vienna 1

All structures that were modified in the Eterna100-V2 for solvability in Vienna 2 were tested for solvability under Vienna 1 through puzzles on Eterna. We found that all except for puzzle 97, “Hoglafractal”, were solvable in both folding engines. This remaining puzzle appears to be unsolvable due to a single base pair at the beginning of the puzzle which is incompatible with Vienna 1 energetic parameters for internal loops and dangling ends. The otherwise solvability of 99/100 Eterna100-V2 puzzles in both Vienna 1 and Vienna 2 folding engines suggests that the redesigns preserved the structural features that made the original puzzles challenging, and the two benchmark versions remain comparable.

### RNA design algorithm rankings are consistent in Eterna100-V2, but performance changes reflect modeling priors

After players modified the secondary structures for the 19 Eterna100 puzzles, we assessed the performance of six RNA inverse folding algorithms on this updated benchmark with publicly available code (Fig. 4): three algorithms relying on Monte Carlo methods, RNAinverse, DesiRNA and NEMO; and three algorithms that also included deep neural networks to guide search, EternaBrain, SentRNA, and LEARNA (**Fig. 4** and **Table 1**; more detailed results in **Supplemental Data Tables S2** and **S3**).

**Fig 4.**
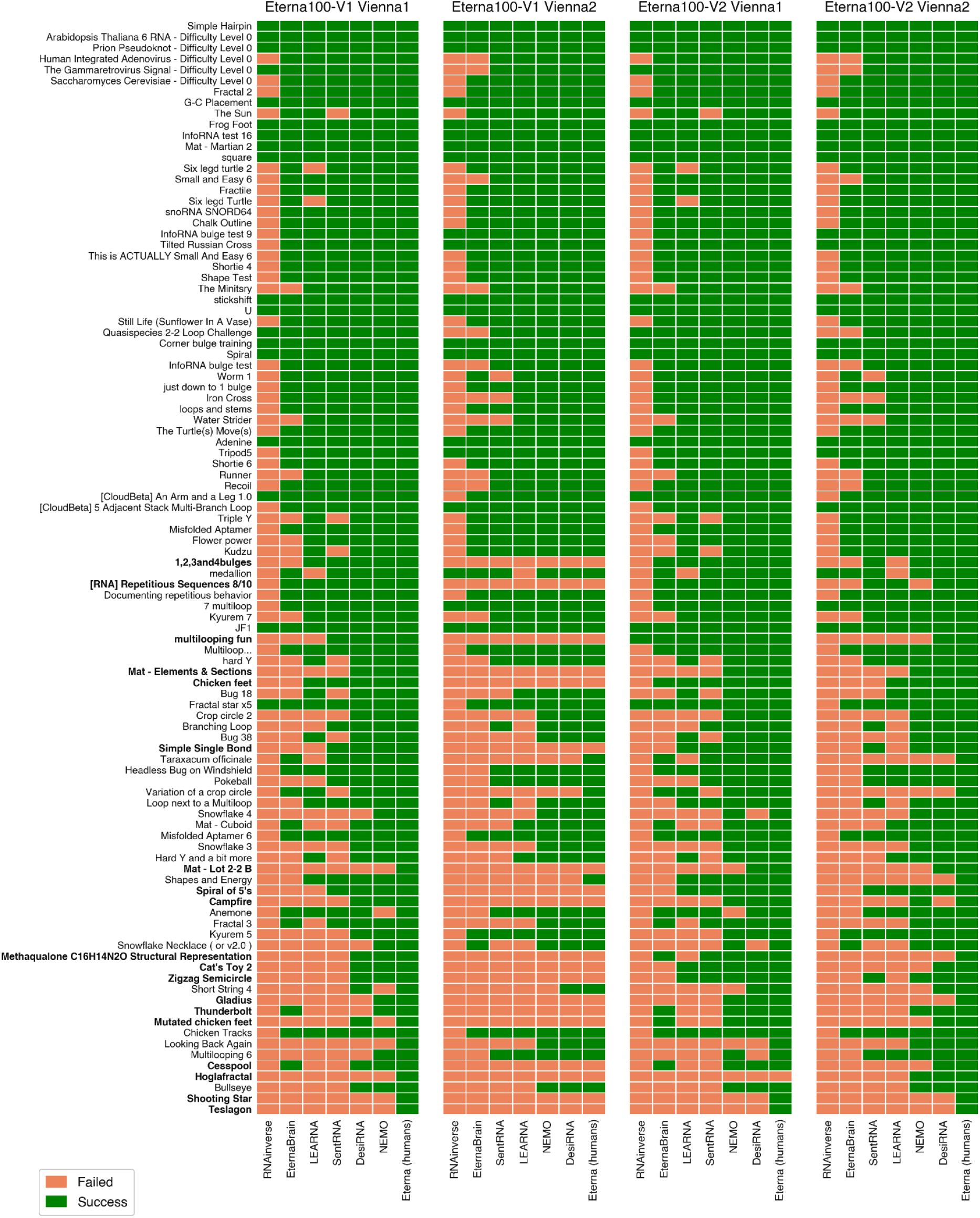
Performance of six algorithms for RNA secondary structure design on the Eterna100-V1 and Eterna100-V2 benchmarks, using either Vienna 1 or Vienna 2 as the folding engine at runtime. Green: Solved; Orange: Unsolved. Bolded names indicate puzzles changed between Eterna100-V1 and Eterna100-V2 benchmarks. Columns are sorted by algorithm performance in each panel.

To evaluate the difficulty of the new Eterna100-V2 benchmark, we tested the algorithms first with the Vienna 1 folding engine, which was also the engine used during development of most of the algorithms. For all of the algorithms, performance on Eterna100-V2 improved compared to Eterna100-V1 when retaining the Vienna 1 folding engine (**Fig. 4** and **Table 1**; RNAinverse, 18 to 19; EternaBrain, 60 to 65; LEARNA, 67 to 72; SentRNA, 69 to 72; DesiRNA, 90 to 93; and NEMO, 92 to 93). This was expected, since the 19 redesigned targets were ‘relaxed’ to make them solvable in Vienna 2, and likely also admit more solutions in Vienna 1 than the originals did. Inspection of particularly difficult puzzles further supports this picture. For example, the 19 structures from Eterna100-V1 that were deemed unsolvable in Vienna 2 were of particular interest (boldface in Fig. 4) as they were some of the hardest secondary structures: in the original Eterna100-V1 benchmark tested in the Vienna 1 folding engine, EternaBrain and SentRNA solved 3 (16%) and 6 (32%) out of the 19, respectively, lower than their average across all Eterna100-V1 puzzles (60% and 69%, respectively). All design methods demonstrated performance improvement in going from the 19 puzzles in Eterna100-V1 to Eterna100-V2 (RNAinverse, 0 to 1; EternaBrain and LEARNA, both 3 to 8; SentRNA, 6 to 9; DesiRNA, 13 to 16; and NEMO, 14 to 15; **Fig. 4** and **Supplemental Data Tables S2 and S3**). Again, this was expected given the increased solution space afforded by the redesigned structures for these 19 puzzles in Eterna100-V2 compared to Eterna100-V1 (for example, through smaller internal loops creating more favorable energetic conditions; **Fig. 1a-b** and **Fig. 3**).

The intended use case for the Eterna100-V2 is to benchmark RNA design algorithms with the more recent Vienna 2 folding engine. When run on the Eterna100-V1 with Vienna 1 (original settings) versus on the Eterna100-V2 with Vienna 2 (modernized settings), performance of the six tested algorithms got better or worse, depending on the algorithm (**Fig. 4** and **Table 1**). RNAinverse solved 18 puzzles under Eterna100-V1 with Vienna 1, which increased to 20 under the Eterna100-V2 with Vienna 2; but EternaBrain’s performance decreased dramatically, from 60 to 49 solved puzzles. LEARNA’s performance increased slightly from 67 to 69, SentRNA decreased slightly from 69 to 68, DesiRNA increased slightly from 90 to 92, and NEMO decreased from 92 to 86. Despite these changes, the rankings of the algorithms were similar in Eterna100-V2 with Vienna 2 compared to Eterna100-V1 with Vienna 1. LEARNA and SentRNA switched places, but they were effectively tied in both benchmarks, with the number of solved puzzles by the two algorithms within 2. In addition, DesiRNA became the top performer in Eterna100-V2 with Vienna 2, solving 6 more puzzles than NEMO, which was the top method for Eterna100-V1 with Vienna 1. There was no clear trend in whether puzzles became more or less difficult in the modernized Eterna100-V2 with Vienna 2 vs. the original Eterna100-V1 with Vienna 1 – some puzzles were solved by fewer algorithms and some by more algorithms (**Fig. 5**). Overall, however, the 19 puzzles that were modernized in Eterna100-V2 to be solvable in Vienna 2 did appear to be the hardest (**Supplemental Fig. S2** and **Supplemental Data Table S4**). These were the puzzles whose original targets proved unsolvable under Vienna 2, and their redesigns were selected to preserve the unique structural elements of the originals (**Fig. 1c**), so their continued difficulty was expected.

**Fig 5.**
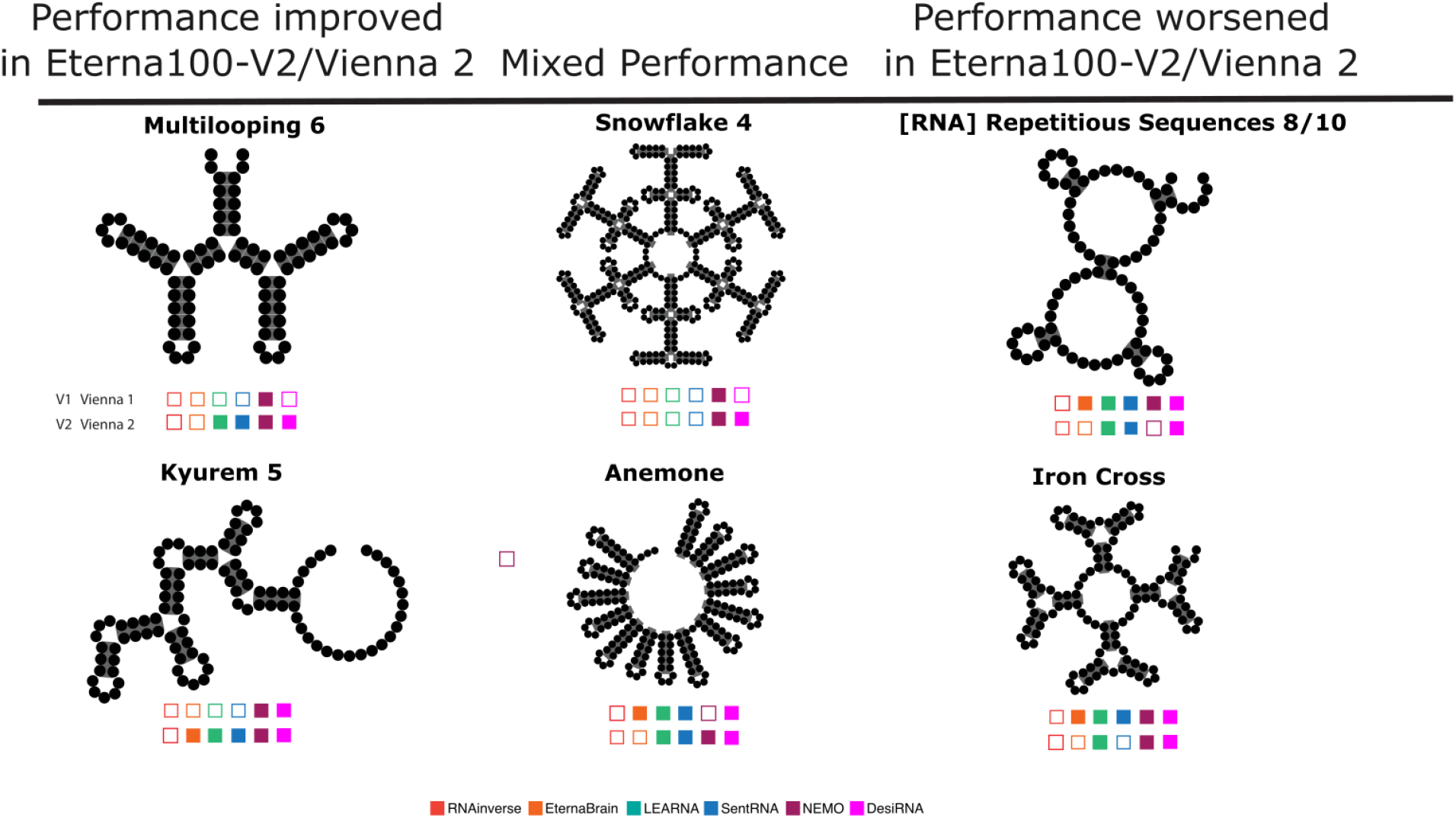
Selected puzzles that demonstrate differences in algorithms’ puzzle-solving ability. Open squares indicate that a given algorithm was unable to solve that puzzle; filled squares indicate that the algorithm did solve the puzzle. The first row of squares indicates algorithms’ performance on Eterna100-V1 using Vienna 1, and the second row indicates the performance on Eterna100-V2 using Vienna 2.

The three algorithms whose success rates dropped (EternaBrain, SentRNA, NEMO) included heuristics and/or neural network training examples that were derived from Eterna players’ efforts with the original benchmark’s Vienna 1 folding engine, suggesting that those sequence design methods were overly dependent on details of that folding engine. Re-training of SentRNA and EternaBrain neural networks with Vienna 2 instead of their original Vienna 1 folding engine changed the number of solved Eterna100-V2 puzzles by 1 or less (**Supplemental Data Table S2)**, indicating that regenerating the networks’ features under Vienna 2 does not recover performance. Both algorithms also use heuristics to refine neural network solutions; their folding-engine dependence thus seems likely to arise from these heuristics, which were developed through Eterna gameplay under Vienna 1.

Overall, the modernized Eterna100-V2 with Vienna 2 remains unsolved algorithmically. Even if solutions are aggregated across the 6 algorithms, 7 problems remain unsolved in Eterna100-V2 with Vienna 2 (**Table 1**). Further runs with 12 CPUs instead of 1 CPUs and 10 trials instead of 1 trial alloted for DesiRNA design runs still left 4/100 problems unsolved. Studies by the DesiRNA authors, based on an early preprint of this article and the Eterna100-V2, leave 3/100 unsolved [27]. It will be interesting to see whether DesiRNA, Montparnasse, or other emerging methods can achieve solutions to the Eterna100-V2.

### Differences in folding engine settings in prior solutions of the Eterna100-V1

Two recent studies reported solving all 100 Eterna100-V1 puzzles with new algorithms: DesiRNA [27], also tested above, and Montparnasse, a nested rollout policy adaptation search [28,34]. Both emulated the benchmark’s original Vienna 1 folding engine by running the ViennaRNA 2 RNAfold algorithm specifying Turner 1999 parameters. We revisited these studies’ solutions after we could not reproduce DesiRNA’s reported performance on Eterna100-V1 (**Table 1**). Re-scored against the Vienna 1 executable, the published DesiRNA designs fold correctly for only 69 of 100 targets and the Montparnasse designs for only 27 of 100. In both cases, discrepancies with the published results are due to a shift between the default setting of dangling end energies changed in Vienna 2. The “dangles” flag sets the number of unpaired nucleotides adjacent to a base pair that can receive energetic bonuses, and the default has changed from 1 to 2; see **Supplemental Data Table S5** and **Methods**. In addition, for Montparnasse, an acceptance criterion that admitted designs within one base pair of the target resulted in incorrect solutions of the Eterna100 benchmark, which rewards only exact match of the secondary structure [28,34]. Recomputing structures for other algorithms that have made sequences for Eterna100-V1 available (MoiRNAiFold [35], aRNAque [36]) also confirmed that they have not used the intended ‘dangles’ settings when emulating Vienna 1 with Vienna 2 (**Supplemental Data Table S5**). To mitigate future confusion, we give the recommended executables, parameters, settings, and example output in **Table 2** for both Eterna100-V1 and Eterna100-V2, and highlight the issues in the online repository at https://github.com/eternagame/eterna100-benchmarking.

**Table 2.**
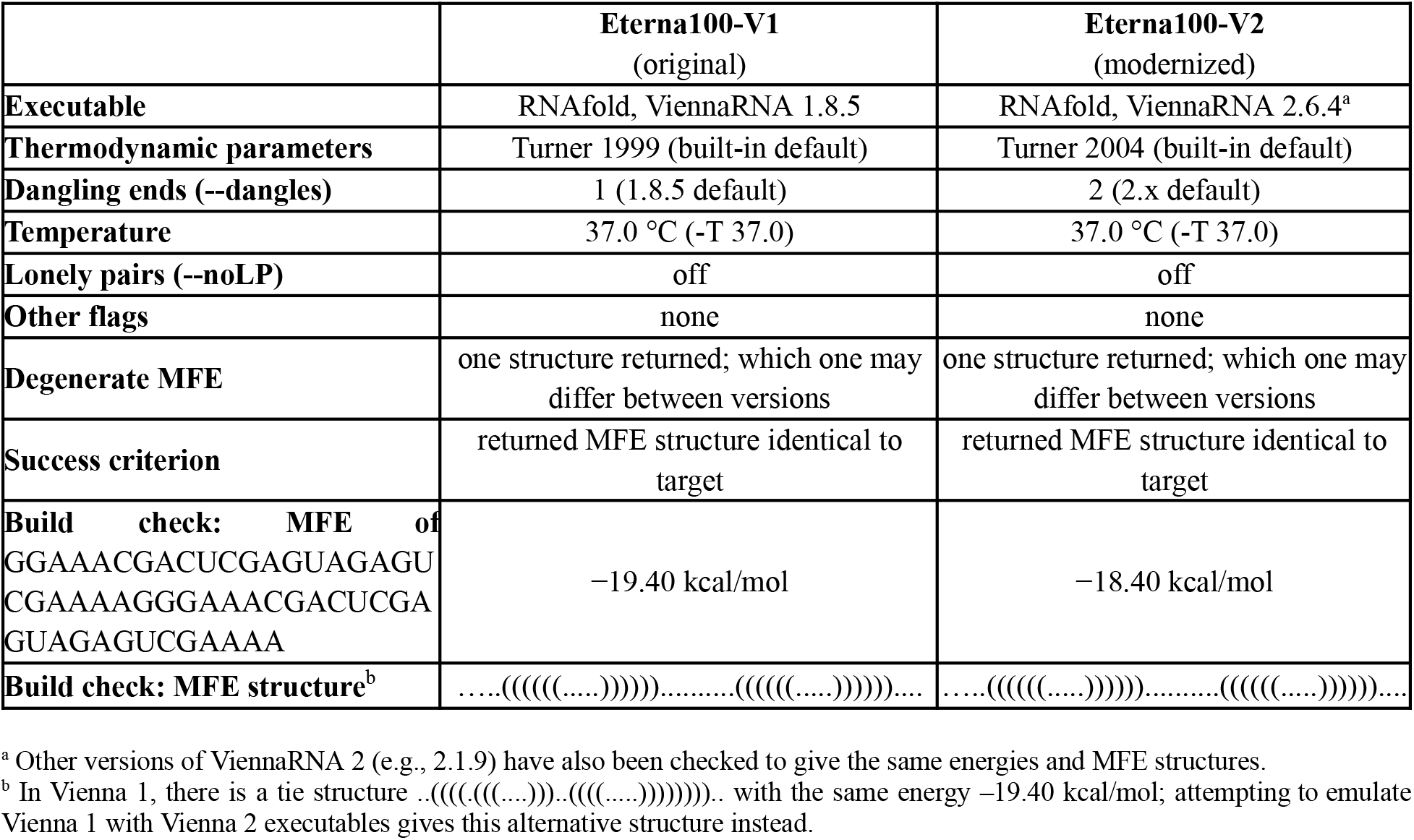
Folding-engine settings defining each Eterna100 benchmark. A build reproducing the build-check rows is correctly configured.

## Discussion

In this work, we have reported that 19 of the 100 structures in the widely used Eterna100 benchmark for evaluating RNA inverse design are unsolvable when the thermodynamic parameters of Vienna 1 were substituted with those of Vienna 2; recent work has supported these observations with rigorous proofs [37]. These results present a problem for evaluating inverse folding algorithms, since their performance depends on the thermodynamic model chosen to develop and to test each algorithm. To amend this problem, we asked Eterna participants to redesign the 19 puzzles. Participants found strategies to do so and proposed redesigned structures that would largely preserve the original challenges of the puzzles. The final new benchmark, termed Eterna100-V2, has 100 of 100 puzzles solvable in Vienna 2.

We next evaluated state-of-the-art algorithms on this updated Eterna100-V2 benchmark. We found that changes in performance reflect model priors. For example, EternaBrain, which was developed with player solutions and heuristic strategies generated under Vienna 1, exhibited notably worse performance with the Vienna 2 folding engine. Additional tests were performed to retrain neural network components of the design methods using folding engines “matching” the respective benchmark but identified minimal impact on original model quality. Instead, other components of the EternaBrain and NEMO models besides their neural networks, such as heuristic design rules based on Vienna 1 gameplay, may reduce their ability to generalize to the Vienna 2 folding engine. In contrast, the newer Monte Carlo method DesiRNA shows improved performance in the Eterna100-V2 benchmark, though it still cannot find solutions to all 100 problems. Motivated by a mismatch in our results on the Eterna100-V1 compared to performance summaries from algorithms like DesiRNA, we also revisited reported solutions to Eterna100-V1 and discovered that settings used to emulate Vienna 1 parameters in the Vienna 2 package led to incorrect solutions.

Given the number of secondary structures that were unsolvable in the original Eterna100 benchmark with the Vienna 2 parameters, it seems likely this benchmark will need to be continuously updated as RNA structure prediction becomes more accurate. Folding engines like CONTRAfold [38] and EternaFold [39] demonstrate improved performance over Vienna folding engines in predicting experimental thermodynamic ensembles and generally predict less stable structures than the Vienna folding engines for the same sequences. Furthermore, metrics beyond minimum free energy structure are potentially better suited [31] for predicting *in vitro* and *in vivo* structural viability, in addition to other desired characteristics which are not reflected by the dominant folded structure alone, such as chemical stability and, for mRNA, protein expression [40].

When solved by players online, Eterna puzzles add constraints beyond the target structure and folding engine. Constraints include nucleotides locked to a given identity at given positions, which are central to the NP-completeness proof for RNA secondary structure design [41], and bounds on GC, GU and AU pair counts or base composition. Players have solved all of Eterna100-V1 and Eterna100-V2 under these constraints; as more advanced challenges for algorithms, these constraints are now provided with validator scripts in the eterna100-benchmarking GitHub repository (“eterna100_player_constraints.csv”) and example solutions satisfying constraints are given there as well as in **Supplemental Data Table S1**. Eterna solutions can also be degenerate, with the displayed MFE structure tied in energy against other structures and the tie broken by package-specific heuristics (**Table 2**); enforcing a unique MFE structure would be stricter [41], but it is unclear if all Eterna100 puzzles admit such a non-degenerate solution.

Given these variations, future RNA inverse folding algorithms should strive to be folding-engine independent, or at least rapidly retrainable with new folding engines and specialized “win conditions”. We hope that open availability of Eterna100-V1 and Eterna100-V2, and further updates from the Eterna community, will enable the continual testing of such algorithms until Eterna is “solved”.

## Methods

### Design challenges on Eterna platform

Through iterative manual design, we identified 19 secondary structures in the original Eterna100 (Eterna100-V1) benchmark that we hypothesized were not solvable using Vienna 2. We asked the Eterna community to redesign these 19 puzzles to be compliant with the thermodynamic parameters of the ViennaRNA 2.1.9 software package as implemented in Eterna (solutions were checked to be valid in later versions of the Vienna RNA 2 code up to Vienna 2.7.1). Players achieved this redesign through the Eterna “Puzzlemaker” interface, whereby individual base pairs and bases may be deleted or added to a structure. Players submitted 52 total puzzles as modifications of the unsolvable 19. Nineteen of those submissions were chosen to both maintain the constraints of the Eterna platform and the identity of unique structural elements that existed in the original puzzles.

During this process, we also discovered that for one of the puzzles (puzzle 5, The Gammaretrovirus Signal - Difficulty Level 0) in the Eterna100-V1, the structure did not match that of the Eterna puzzle it was derived from (and so neither did the sample solutions solve the structure). This was due to an error which caused a single nucleotide to be omitted - we have treated the Eterna100-V1 structure as correct and provided an updated Eterna puzzle ID and sample sequence for Vienna 1 in **Supplemental Data Table S1**.

### Automated tests of RNA secondary structure design algorithms

We evaluated the performance of six algorithms (EternaBrain, SentRNA, RNAinverse, LEARNA, NEMO, and DesiRNA) on the Eterna100-V1 and on the Eterna100-V2 with the 19 modified puzzles, using ViennaRNA 1.8.5 as well as the more recent ViennaRNA 2.6.4 as the folding engine. Separate runs were performed for each puzzle, algorithm, and Vienna version combination (with the exclusion of the 19 unsolvable Eterna100-V1 puzzles and one unsolvable Eterna100-V2 puzzle under Vienna1, on the basis of the Eterna participants’ failed attempts and the extended NEMO search described below). Algorithms were run for a maximum of 24 hours with a single AMD 7502 core on the Stanford University Sherlock cluster. We note that use of the ViennaRNA 2 package with Turner99 parameters, as carried out recently [27], can be used to emulate ViennaRNA 1 but must use the setting “--dangles=1” to reproduce the behavior of the Vienna 1 folding engine. Additionally, where multiple structures have the same free energy for a given sequence, Vienna 2 run in this way may select a different structure as the MFE structure than Vienna 1, resulting in a sequence that seems to solve a puzzle actually failing with the intended folding engine (see also **Table 2**).

EternaBrain uses a combination of a convolutional neural network trained on Eterna player moves and a Single Action Playout (SAP) [23], a depth-1 Monte Carlo Search using Eterna player strategies. In addition to the pretrained CNN trained with features generated by Vienna 1, the CNN was retrained using features generated by Vienna 1 and Vienna 2. Variants were tested using combinations of the Vienna 1 or Vienna 2 trained CNN, using Vienna 1 or Vienna 2 during inference to generate features passed to the CNN, and using either Vienna 1 or Vienna 2 during the SAP step. Variants where the folding engine in the SAP step was different from the engine used to generate features for the CNN inference step (termed “flipped”, eg vienna1-flipsap run in “vienna1” mode uses the CNN trained using Vienna 1 features, with Vienna 1 for the CNN inference features, and Vienna 2 for the SAP) were tested as the player moves used to train the CNN were all using Vienna 1 originally, and insufficient data exist to train a neural network with Vienna 2 player moves. The “extended” 92-puzzle dataset from the latest version of the EternaBrain repository was used, rather than the published 78-puzzle dataset. Training was run until completion.

SentRNA, unlike EternaBrain, does not rely on player moves to train its deep neural networks; instead, the authors define their own set of features in the Methods section of the paper (though it does rely on player solutions generated under Vienna 1) [24]. In addition to using the pretrained models (selecting the 20 models trained using 20 structural features), under both Vienna 1.8.5 and 2.6.4, we re-trained an ensemble of 20 networks (each using 20 structural features) with 300 “solution trajectories” [24]. Training was run until completion. For LEARNA, in addition to the pretrained model, we attempted to similarly retrain the reinforcement learning models using Vienna 1.8.5 and 2.6.4, keeping the original hyperparameter and architecture values (i.e., without performing a new joint architecture and hyperparameter search). Training was run for one hour on 20 CPUs on the Stanford University Sherlock cluster, but did not lead to performance competitive with the pre-trained LEARNA models made available with the publication. DesiRNA was benchmarked here by us rather than taken from its publication. Because the released code exposes no dangling-end option, we patched it to add a --dangles flag, and ran it with --dangles=1 for Vienna 1 and --param=2004 for Vienna 2. (As noted above, for Vienna 1, loading the Turner 1999 parameters alone leaves dangles at the ViennaRNA 2 default of 2, which reproduced neither folding engine.) DesiRNA solutions for Vienna 1 settings were confirmed to be solutions with the Vienna 1.8.5 executable. DesiRNA was given the same budget as other algorithms reported here, one trial per puzzle on a single CPU with a 24-hour cap. We note that a larger budget of 12 CPUs and 10 independent trials per puzzle, with the same wall-clock cap, raised DesiRNA from 92 to 96 of 100 on Eterna100-V2 with Vienna 2, and from 90 to 92 of 100 on Eterna100-V1 with Vienna 1.

RNAinverse and NEMO, which are both stochastic methods, were able to be run without modification using either Vienna 1 or Vienna 2 as the internal folding engine. RNAinverse was benchmarked using standard settings, allowing either Vienna 1.8.5 or Vienna 2.6.4 to be used as the folding algorithm. NEMO was run with a maximum of 2500 Monte Carlo iterations. For Vienna 1 settings, NEMO, like DesiRNA, uses library calls into the Vienna 2 package but with Turner1999 parameters and dangles=1; NEMO solutions under these settings were confirmed to be solutions with the Vienna 1.8.5 executable. To more stringently test the unsolvability of the 19 “unsolvable” Eterna100-V1 puzzles in Vienna 2, NEMO was given 10 independent 24-hour periods to attempt to solve using 16 (unspecified) cores with a maximum of 10^4^ Monte Carlo iterations. No solutions were found.

### Re-scoring of published Eterna100-V1 solutions

Sequences reported as solutions to the Eterna100-V1 were collected from their original sources: the DesiRNA repository listing accompanying Wirecki et al. [27]; Appendices B and C of Cazenave [34], which give the Montparnasse designs and a re-run of DesiRNA, respectively; section C (“EteRNA Sequences”) of the MoiRNAiFold supplementary information [35]; and the two aRNAque mutation variants [36]. Each sequence was folded with RNAfold -T 37.0 from stock builds of ViennaRNA 1.8.5, 2.1.9 and 2.6.4, with no -d, -P or --noLP flags, which is the criterion the benchmark itself applies **(Table 2)**: a design counted as a solution only if the returned minimum-free-energy structure was exactly equal to the target structure. As a positive control, Eterna participants’ own sample solutions scored correctly in each of the four target/engine combinations, confirming that the scoring procedure reproduced the benchmark. To test whether any single model setting reproduced each published listing, every solution set was also scored at all 16 points of the grid formed by the Turner 1999 and Turner 2004 parameter sets, dangling-end treatments 0 through 3, and --noLP on or off. Counts for every solution set at different settings are given in **Supplemental Data Table S5**.

### Use of AI LLMs

Manuscript revision, solution re-scoring, and cross-validation analyses reported here were carried out with the assistance of Claude (Anthropic, Opus 5.5); all reported numbers and conclusions were confirmed by the authors.

## Data availability

Eterna100 sequences, solutions, and validator scripts are publicly available at https://github.com/eternagame/eterna100-benchmarking and https://github.com/eternagame/eterna-validator.

## Acknowledgements

We dedicate this paper to the memory of D. S. Kim. We thank T. Zhao and L. Huang (Oregon State), J. Bujnicki and lab (IIMCB, Warsaw), T. Castenave (U. Paris, Dauphine), A. Panescu

(Yale), and A. Kohli (U.C. Berkeley) for correspondence on Eterna100 solvability. Calculations and model training were performed on the Stanford Sherlock cluster and Berkeley CSUA cluster. We acknowledge funding from the National Science Foundation (GRFP to H.K.W.S.), the National Institutes of Health (R35 GM122579 to R.D.), and the Howard Hughes Medical Institute (R.D.).

## Author Contributions

B.R. and R.D. formulated the problem. R.V.K., J.R., and H.K.W.S. evaluated RNA design algorithms. R.V.K., B.R., and J.R. performed analyses. Eterna Participants designed the secondary structures and generated additional solutions, as follows. B.R. (Brourd), Tesla’sDisciple, W.A.D. (Astromon), c-quence, JR, Jieux, jnicol, clollin created the updated 19 puzzles in Eterna100-V2 with minimal changes. wawan151, Jieux, lroppy (Leonard Oppenheimer), jnicol (John Nicol), and JSci solved puzzles without algorithmic solutions. W.A.D. performed checks on solutions, including solvability of Eterna100-V2. R.V.K., B.R., J.R., H.K.W.S., and R.D. wrote the manuscript.

## Notes

https://github.com/eternagame/eterna100-benchmarking

https://github.com/eternagame/eterna-validator

### Competing Interest Statement

The authors have declared no competing interest.

### Summary of Updates

Completion of benchmarking, including DesiRNA runs; checks of validity of published Montparnasse and DesiRNA solutions; explicit recommendations for reproducing intended Vienna 1 & Vienna 2 settings; example solutions that obey additional constraints imposed for game players; dedication to Do Soon Kim.

