## Supplementary Material for "Redesigning the Eterna100 RNA design benchmark for the Vienna 2 folding engine"

**Supplementary Materials:**

- **Supplementary Figures** S1-S2 (this document)
- **Supplementary Data** (Tables S1-S5), attached separately in a single Excel file

ANNOUNCEMENTS

30 Jan 2020

### Eterna100-V2: Impossible(?) puzzles and call for solutions

Hi Eterna Players,

A few months ago, in the creation of the Eterna100-V2, we asked that players design a number of secondary structures modifications to 19 puzzles in the Eterna100 that made them solvable in the Vienna2 energy model as implemented on Eterna. However, the question as to whether these puzzles were truly unsolvable in these parameter sets was never really put forth towards players. As some of you may have recently seen, there is a new series of puzzles that have been published on the site that try to address this question. The [IMOSSIBLE?] puzzles are those original 19 Eterna100 puzzles that we believe are unsolvable in Vienna2, published in Vienna2. However, determining if a secondary structure is unsolvable is a hard problem, and there is a possibility that we missed potential solutions!

In addition to those puzzles that we believe are unsolvable, there are also the Vienna2 remakes of the Eterna 100 puzzles. Referred to as Eterna100-v2, we are publishing them in Vienna1.8.5 to determine if these structures are backwards compatible in their energy functions. Some of these we have solutions for, but others we are not so sure about!

So we ask that Eterna players try solving these puzzles, and if you can't solve them that's okay. In the comments of the puzzle, you can post which base pairs you were not able to stabilize, and that could provide us with greater insight on what makes these puzzles unsolvable. Below are links to the lists of puzzles that we published on the site.

[IMPOSSIBLE?][Vienna 2][Eterna100-v1]
[Eterna100-v2][Vienna 1]

**Supplemental Fig. S1.** Eterna announcement asking players to attempt to solve the nineteen modified puzzles in Vienna 1 and Vienna 2.

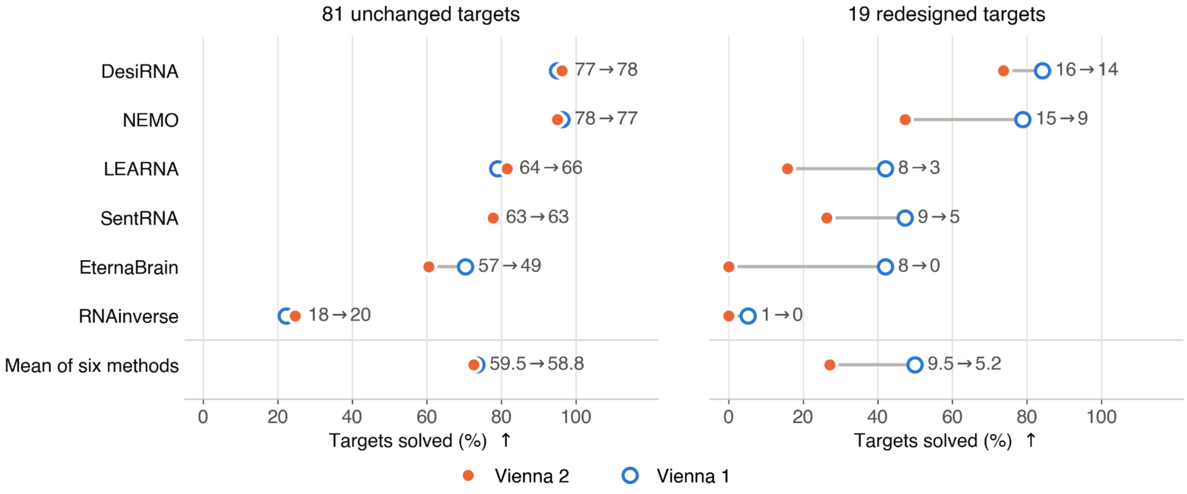


a

**Supplemental Fig. S2. Targets re-designed for Eterna100-V2 are more difficult in Vienna 2 than Vienna 1.** The same 100 Eterna100-V2 targets scored with Vienna 1 and with Vienna 2, split into the 81 carried over unchanged from Eterna100-V1 and the 19 redesigned by Eterna participants. Although 'relaxed' to enable solution in Vienna 2, the redesigned targets sit close to the edge of what is designable in Vienna 2.
